# Distinct accessory roles of Arabidopsis VEL proteins in Polycomb silencing

**DOI:** 10.1101/2023.05.22.541744

**Authors:** Elsa Franco-Echevarría, Mathias Nielsen, Anna Schulten, Jitender Cheema, Tomos E Morgan, Mariann Bienz, Caroline Dean

## Abstract

Polycomb Repressive Complex 2 (PRC2) mediates epigenetic silencing of target genes in animals and plants. In Arabidopsis, PRC2 is required for the cold-induced epigenetic silencing of the *FLC* floral repressor locus to align flowering with spring. During this process, PRC2 relies on VEL accessory factors, including the constitutively expressed VRN5 and the cold-induced VIN3. The VEL proteins are physically associated with PRC2, but their individual functions remain unclear. Here, we show an intimate association between recombinant VRN5 and multiple components within a reconstituted PRC2, dependent on a compact conformation of VRN5 central domains. Key residues mediating this compact conformation are conserved amongst VRN5 orthologs across the plant kingdom. By contrast, VIN3 interacts with VAL1, a transcriptional repressor that binds directly to *FLC*. These associations differentially affect their role in H3K27me deposition: both proteins are required for H3K27me3, but only VRN5 is necessary for H3K27me2. Although originally defined as vernalization regulators, VIN3 and VRN5 co-associate with many targets in the Arabidopsis genome that are modified with H3K27me3. Our work, therefore, reveals the distinct accessory roles for VEL proteins in conferring cold-induced silencing on *FLC*, with broad relevance for PRC2 targets generally.

## Introduction

Polycomb Repressive Complex 2 (PRC2) is pivotal for the transcriptional silencing of genes that control development and differentiation in animals and plants. Silencing consists of an epigenetic memory mechanism based on the deposition of trimethylated lysine 27 of histone H3 (H3K27me3) at Polycomb target genes (Cao et al. 2002; Czermin et al. 2002; Muller et al. 2002). PRC2 comprises four subunits, each deeply conserved throughout animals and plants (Birve et al. 2001; Tie et al. 2001) (**Supplemental Fig. S1A**). These cooperate to deposit the H3K27me3 silencing hallmark at cis-regulatory regions of targets and to copy it to daughter strands during DNA replication in a ‘read-write’ mechanism (Yu et al. 2019). PRC2 accessory proteins are also required for long-term epigenetic memory when cell division rates are high (Yu et al. 2019; Lovkvist et al. 2021; Menon et al. 2021). They have been found to modulate the methyltransferase activity of PRC2 and/or promote tethering of the complex at genomic loci (Hauri et al. 2016; Choi et al. 2017; Chen et al. 2018; Kasinath et al. 2018; Oksuz et al. 2018; Hojfeldt et al. 2019; van Mierlo et al. 2019).

In Arabidopsis and other plants, some of the PRC2 subunit functions are encoded by multiple genes, for example, the SUZ12 function can be provided by VRN2, EMF2 or FIS2, and these are expressed in different developmental contexts (Mozgova and Hennig 2015; Godwin and Farrona 2022). The best-known complex is VRN2-PRC2 whose function has been extensively characterised in vernalization, i.e. the cold-induced epigenetic silencing of the *FLOWERING LOCUS C* (*FLC*) (Michaels and Amasino 1999; Hepworth and Dean 2015; Xu and Chong 2018; Costa and Dean 2019; Sharma et al. 2020). In contrast to the deep conservation of the PRC2 subunits (**Supplemental Fig. S1A**), the accessory proteins vary between animals and plants, often without obvious counterparts in the other kingdom. Amongst the best-known plant accessory factors are the VEL proteins: *VIN3* and *VRN5* were identified genetically in Arabidopsis as genes required for *FLC* silencing (Sung and Amasino 2004; Greb et al. 2007). VRN5 and VIN3 physically associate with PRC2 in cold-treated plants along with the third homologue VEL1 (De Lucia et al. 2008; Kim and Sung 2010).

The hallmarks of VEL proteins are an atypical plant homeodomain (PHD) zinc finger, a fibronectin type III (FN3) domain and a C-terminal VEL domain (Sung and Amasino 2004; Wood et al. 2006; Greb et al. 2007) (**Fig. 1A**). We previously discovered that the VEL domain engages in spontaneous head-to-tail polymerization to assemble biologically relevant, dynamic biomolecular condensates (Fiedler et al. 2022). We have also shown that the PHD finger is the core module of a compact tripartite PHD superdomain (designated PHDsuper below) without histone H3 tail binding activity (Franco-Echevarría et al. 2022). In Arabidopsis plants, VIN3 expression is cold-inducible while VRN5 is constitutively expressed, and both become enriched at the Polycomb nucleation region of *FLC* during prolonged cold exposure (Sung and Amasino 2004; Yang et al. 2017). Their association with the *FLC* nucleation region in the cold confers metastable silencing, with the H3K27me3 spreading across the whole locus upon return to warm temperatures to confer long-term epigenetic stability (Yang et al. 2017).

**Figure 1.**
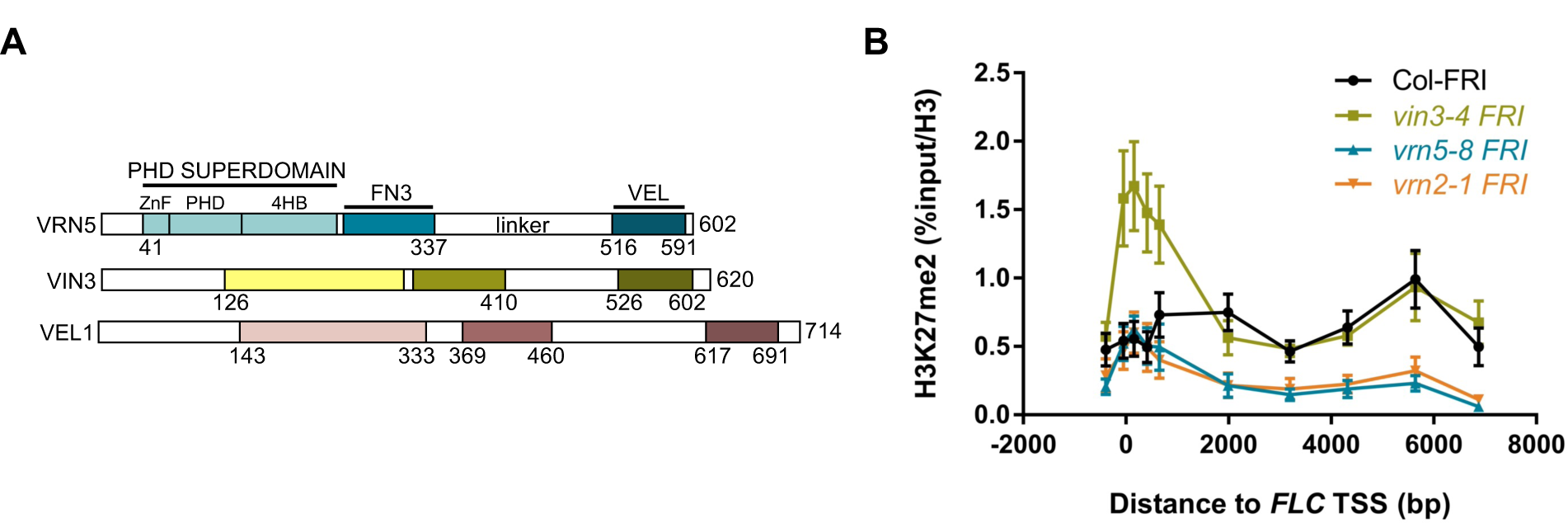
H3K27me2 enrichment at *FLC* in wt and mutant Arabidopsis. (A) Domain organization of VEL proteins. (B) Enrichment of H3K27me2 levels at *FLC* in Col-FRI and different mutant plants vernalized for 6 weeks (6WT0). Data are shown as the percentage input relative to H3. Non-transgenic Col-FRI plants were used as a control sample. Error bars are means ± s.e.m. from two independent experiments.

Recent studies have identified VEL relatives in other plants that do not undergo vernalization (Mohan et al. 2018; Yang et al. 2019; Fiedler et al. 2022; Shwartz et al. 2022) raising questions about the generality of VEL protein function and how they connect with PRC2 silencing. We, therefore, investigated the individual roles of the VEL proteins in PRC2 silencing, detailed VEL protein structural characteristics that lead to specific functionalities, and mapped the sites in the Arabidopsis genome where they are stably recruited. Our work reveals highly conserved features of different VEL paralogs that mediate differential interactions with PRC2 and other silencing proteins. The genome-wide co-localization of the VEL proteins at numerous other H3K27me3-enriched targets also suggests VEL proteins have a broad role in PRC2 silencing during plant growth and development.

## Results and Discussion

### Differential effects of vrn5 and vin3 on H3K27me2 dynamics

Methylation of H3K27 by PRC2 is intrinsically slow, so that H3K27me3 levels never reach their maximum before cell division occurs (Alabert et al. 2015; Berry et al. 2017). Therefore, one proposed role of the PRC2 accessory proteins is to increase the residency time of PRC2 at targets, consistent with both *vin3* and *vrn5* mutants accumulating no H3K27me3 at the *FLC* nucleation region in the cold. To explore this, we investigated the intermediate methylation state, H3K27me2, at *FLC*. ChIP-qPCR showed no H3K27me2 enrichment in the *FLC* nucleation region in vernalized *vrn5* mutants (6WT0), with *vrn2* (defective in the PRC2 SUZ12 subunit) used as the benchmark control (**Fig. 1B**). Thus, VRN5 and VRN2 are strictly required for the PRC2-dependent deposition of H3K27me2 at *FLC*. Somewhat surprisingly though, *vin3* mutants showed accumulation of H3K27me2 in the nucleation region (**Fig. 1B**), revealing different roles for VIN3 and VRN5 in the PRC2-dependent silencing of *FLC*.

### VRN5 associates tightly with PRC2

Previous proteomic evidence indicated close association between VEL proteins and PRC2 (De Lucia et al. 2008; Questa et al. 2016), but whether these were direct was unknown. To address this, we co-expressed HA-tagged versions of all four PRC2 subunits (VRN2, SWN, FIE and MSI1; **Supplemental Fig. S1B**) with GFP-tagged VEL proteins in heterologous human embryonic kidney (HEK293T) cells to avoid potential bridging interactions from endogenous proteins (Fiedler et al. 2022); and monitored their associations by coimmunoprecipitation (coIP). We found that all four PRC2 subunits coIP efficiently and robustly with GFP-VRN5 but detected much less robust coIP of any PRC2 subunits with VIN3-GFP or GFP-VEL1 (except for a slightly weaker coIP of MSI1 with the latter; **Fig. 2A**). This identifies VRN5 as the main and direct interaction partner of PRC2.

**Figure 2.**
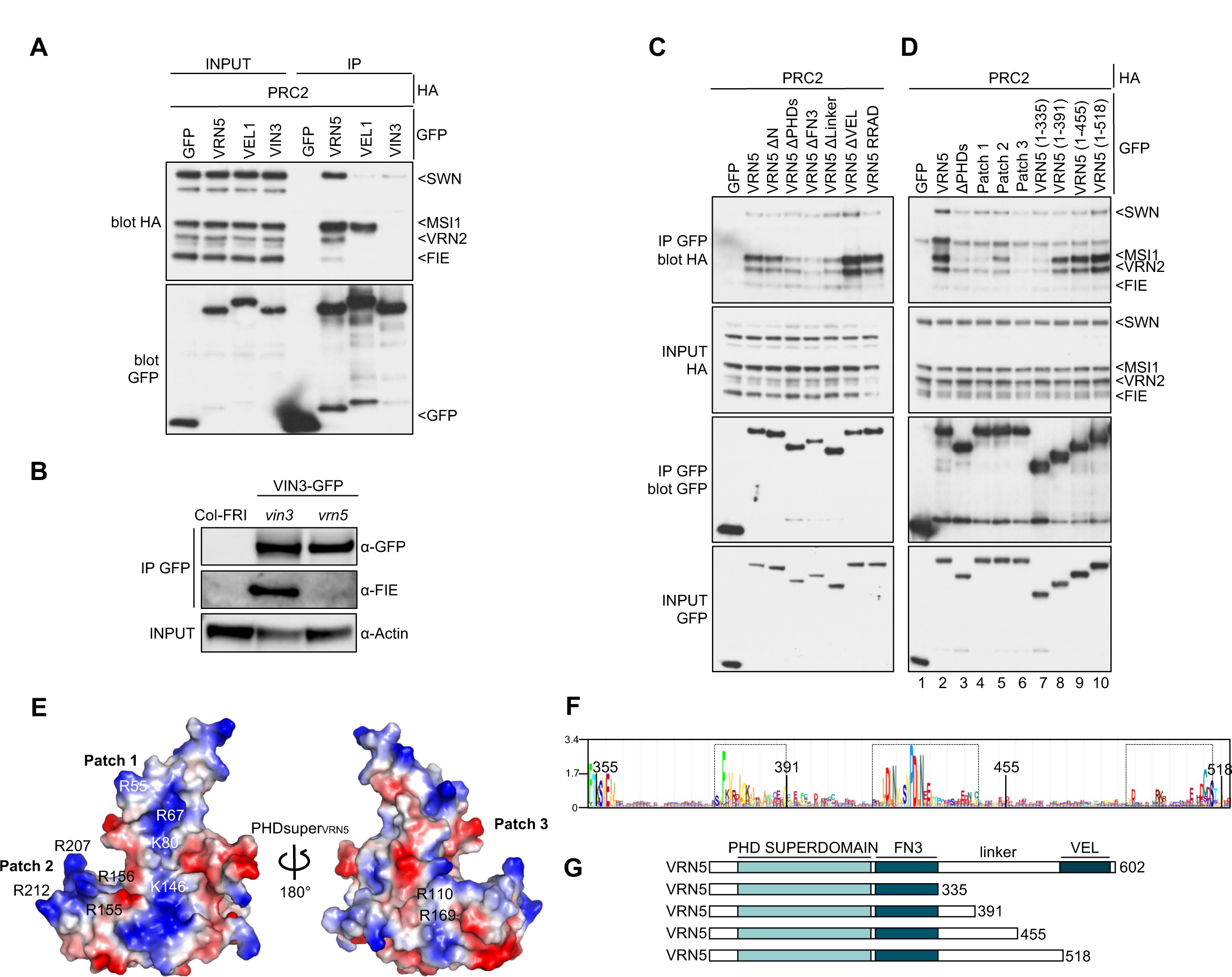
Analysis of the VRN5-PRC2 interaction. (A) coIP of HA-tagged PRC2 core components with GFP-tagged VEL proteins, following co-expression in HEK293T cells, as indicated in panels. (B) Western blots of a-GFP immunoprecipitates from extracts of vernalized *vrn5* mutant plants bearing a *VIN3-GFP* transgene, probed with a-FIE antibody. (C) coIP of HA-tagged PRC2 core components with wt or mutant GFP-VRN5 bearing internal domain deletions, as in (A). (D) coIP of HA-tagged PRC2 core components with wt or GFP-VRN5 mutants as described in (G). (E) Molecular surface representation of PHD_VRN5_ predicted by AlphaFold2, coloured according to electrostatic potential (*red*, negative; *blue*, positive), showing conserved positively charged residues forming cluster 1 and 2 (*front surface*) and cluster 3 (*rear surface*). (F) Sequence logo conservation analysis of VRN5 linker connecting FN3 and VEL domains in viridiplantae, with residue numbers in panel; *dashed squares*, three conserved regions (residues 371-391, 420-445 and 493-518 in Arabidopsis). (G) Schematic representation of different VRN5 linker constructs used in (D).

The lack of a robust coIP between VIN3 and PRC2 was surprising since previous coIP assays in plants revealed that VIN3 co-purified with VRN2 in extracts from vernalized plants and co-fractionated with PRC2 subunits during size exclusion chromatography (Wood et al. 2006; De Lucia et al. 2008). However, these apparent associations might have been bridged by endogenous VRN5. To test this, we probed VIN3-GFP immunoprecipitates from extracts of vernalized *vrn5* mutant plants for the presence of the PRC2 subunit FIE. We thus confirmed that FIE co-purifies with VIN3 in a *vin3* mutant control line (pVIN3:VIN3-GFP in *vin3*), but that it failed to do so in a *vrn5* mutant line (**Fig. 2B**). This is consistent with our coIP results from HEK293T cells **(Fig. 2A)** and indicates that VIN3 is not a direct interactor of the PRC2 core complex. It further suggests that the above-mentioned associations between VIN3 and PRC2 are likely mediated by VRN5, possibly by direct binding between the VEL domains of VIN3 and VRN5 (Greb et al. 2007; Fiedler et al. 2022).

To further define the binding of VRN5 to PRC2, we tested individual GFP-VRN5 deletion mutants for their ability to coIP PRC2. We thus found that PHDsuper, FN3 and the flexible linker between these domains are each required for PRC2 association whereas VEL is not, nor is its ability to polymerize (Fiedler et al. 2022) (**Fig. 2C**). PHDsuper exhibits three surface patches formed by clusters of arginines (R) and lysines (K) that are not seen in canonical PHD fingers (Franco-Echevarría et al. 2022), which bind to positively-charged histone H3 tails (Musselman and Kutateladze 2011; Sanchez and Zhou 2011). Substitutions of these positively charged R and K residues with glutamic acid (E) reduced (in the case of patch 2) or almost abolished (in the case of patch 1 or 3) PRC2 coIP with GFP-VRN5, whereby the effects of patch1 or 3 substitutions were as strong as that of deleting PHDsuper (**Fig. 2D**, lanes 1-6). Therefore, these positively-charged surface patches, predicted to be located in the front and back faces of PHDsuper by Alphafold2 (Jumper et al. 2021) (**Fig. 2E**), are important for the binding of VRN5 to PRC2. Crucially, the equivalent mutants of palm tree VIN3 PHDsuper are soluble, stable, and folded correctly, which indicates that the effects of these mutants in reducing PRC2 association reflect specific interaction defects. Note that patches 1 and 2 are near-invariant amongst plant VEL proteins (Franco-Echevarría et al. 2022) (**Supplemental Figs. S2, S3**), arguing for a wide-spread physiological relevance of these unusual positive surface patches of VEL PHD superdomains.

The requirement of the linker between VRN5 FN3 and VEL for the VRN5-PRC2 association was somewhat unexpected as this linker (comprising 180 residues) is poorly conserved among VEL proteins (**Supplemental Figs. S2, S3**) and predicted to be intrinsically disordered. A search for conserved residues within this linker using HMMER (Potter et al. 2018) identified three short conserved motifs (spanning residues 371-391, 420-445 and 493-518, respectively) within Arabidopsis VRN5 (**Fig. 2F**). To address whether any of these are required for the VRN5-PRC2 interaction, we tested progressive internal deletions retaining increasing linker length (**Fig. 2G**) by coIP and found that the efficiency of PRC2 association with GFP-VRN5 increased with progressively longer linker sequences (**Fig. 2D**, lanes 7-10). This suggests that this linker of VRN5 assists its binding to PRC2, e.g. by wrapping around the PRC2 complex and holding it together. We also found that the N-terminus of VRN5 (predicted to be structurally disordered) contributes to its association with SWN, MSI1 and VRN2, while its C-terminal 11 residues contributes to its association with SWN (**Supplemental Fig. S1C**), indicating that these additional flanking regions assist VRN5 in its binding to PRC2.

Finally, we also confirmed that the plant PRC2 complex, like its human counterpart (Kasinath et al. 2018), is capable of self-assembly, given that SWN, MSI1 and FIE each associate with the GFP-tagged VRN2 (SUZ12) scaffold. Interestingly, if VRN5 is co-expressed with these PRC2 subunits, coIP of MSI1 with GFP-VRN2 increases in a VRN5-dependent manner (**Supplemental Fig. S1D**), suggesting that MSI1 contributes to at least one of the PRC2 docking points for VRN5.

### Different VEL orthologs are evolutionarily conserved throughout the plant kingdom

We have previously shown that PHDsuper and VEL domains are highly conserved throughout the plant kingdom (Fiedler et al. 2022; Franco-Echevarría et al. 2022). Exploring the conservation of the corresponding full-length sequences revealed a high degree of conservation across all plant VEL paralogs. However, VRN5 orthologs differ most from other VEL orthologs, as they are distinguishable from the latter by an invariant DLNxxxVPDLN motif in their linker (**Supplemental Fig. S2**, green dashed square). By contrast, there is a high degree of similarity between VIN3 and VEL1 orthologs which together appear to constitute a subclass of VEL proteins (**Supplemental Fig. S3**). Indeed, VIN3 and VEL1 orthologs can only be distinguished from each other in Brassicaceae, but they cannot be differentiated in more primitive angiosperm species even if these contain multiple VIN3/VEL1-like genes (**Supplemental Fig. S3**). Notably, VRN5 and VIN3/VEL1 orthologs are present in both dicots and monocots, including plant species such as palm tree (*Phoenix dactylifera*, Pd) or maize (*Zea mays*, Zm) that do not require winter cold exposure for spring flowering. This suggests that the different VEL orthologs have widespread roles beyond vernalization in PRC2-dependent silencing.

### Understanding the binding preference of PRC2 for VRN5

The striking preference of PRC2 for VRN5 points to a specialized function of VRN5 orthologs. To understand why VRN5 is the only VEL protein capable of interacting with all four core PRC2 subunits, we used Alphafold2 (Jumper et al. 2021) to predict the structures of different VEL paralogs in Arabidopsis. Interestingly, this revealed a major structural difference between VRN5 versus VIN3 or VEL1: PAE plot analysis by AlphaFold2 (https://alphafold.ebi.ac.uk/) suggests a close packing of PHDsuper against FN3 that seems exclusive to VRN5 orthologs, based on sequence conservation (**Fig. 3A,B; Supplemental Figs. S2, S3**). Indeed, VRN5 orthologs are predicted consistently to adopt a compact conformation in a wide range of plants, whereas the conformation of VIN3 and VEL1 orthologs is predicted to be more open (**Supplemental Fig. S4A**). The backbones of the predicted PHDsuper-FN3 structures of VRN5 orthologs exhibit low RMSD and high template-modeling (TM) score values, ranging from 0.81 to 0.94, whereby 1 indicates a perfect match between two structures. Furthermore, the predicted folds for various PHDsuper-FN3 modules closely resemble each other within the VRN5 lineage (**Supplemental Figs. S4, S5A-C**), suggesting that the interface between PHDsuper and FN3 is deeply conserved amongst all angiosperm VRN5 orthologs.

**Figure 3.**
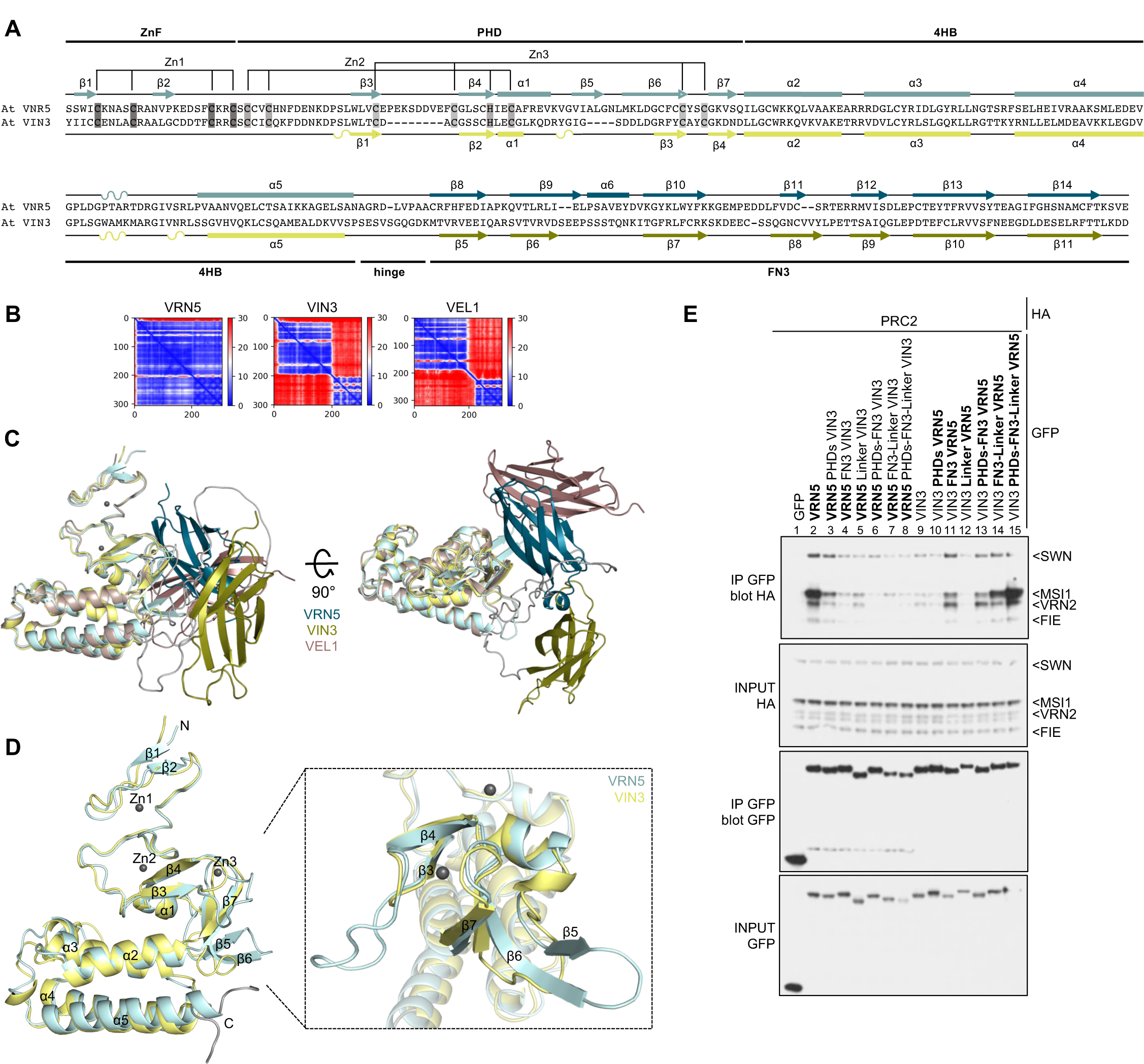
Structural differences between VRN5 and VIN3/VEL1 paralogs. (A) Sequence alignment between At VRN5_41-339_ and At VIN3_126-412_, with predicted secondary structure indicated for VRN5 (*above*) and VIN3 (*below*); highlighted are the Zn^2+^-ligating residues of ZnF and PHD finger. (B) PAE plot obtained for PHDsuper and FN3 domains of At VRN5_41-339_, At VIN3_126-412_ and At VEL1_143-462_ (*blue*, low error; *red*, high error; see also Supplemental Fig. S4). (C) Orthogonal views of superpositions of At VRN5_41-339_ (*blue*), At VIN3_126-412_ (*yellow*) and At VEL1_143-462_ (*brown*) in ribbon representation, as predicted by AlphaFold2; the zinc ions are from Pd PHD_VIN3_ (PDB: 7QCE). (D) Superpositions of PHD superdomains of At VRN5_41-240_ (*light blue*) and VIN3_123-307_ (*light yellow*), as in (C), with secondary structure elements indicated; *grey balls*, zinc ions (RMSD 1.70 Å); close-up view, superimposed proximal PHD fingers as in (D). (E) coIP of HA-tagged PRC2 core components with wt or mutant GFP-VRN5 or GFP-VIN3 bearing internal deletions or domain swaps (indicated above panel), as in Fig. 2A.

Examining the predicted PHDsuper folds of different VEL paralogs, we also found a major difference between VRN5 and VIN3 orthologs: VRN5 exhibits a bulky β-strand extension in the proximal region of the PHD finger, which is missing in VIN3 and VEL1 (**Fig. 3A; Supplemental Figs. S2, S3**). This extension is formed by VRN5 β6 (the topological equivalent to the shorter VIN3/VEL1 β3) which interacts with two shorter adjacent β-strands β7 and β5 in an antiparallel fashion (whereby β5 is unique to VRN5 orthologs; **Fig. 3C,D**). Indeed, the known crystal structure of the PHDsuper from palm tree lacks this β-strand extension (Franco-Echevarría et al. 2022), as do the predicted PHD superdomains from other VIN3 or VEL1 orthologs (**Supplemental Fig. S5D-F**). This distinctive feature characteristic of VRN5 orthologs points to a pivotal role of this structural element in the intramolecular interaction with FN3 (see below).

To test the functional relevance of individual domains for the PRC2-VRN5 interaction, we generated GFP-tagged VRN5 and VIN3 chimeras in which individual domains were substituted with equivalent domains from the opposite type, to monitor their ability to coIP PRC2. We thus found that GFP-VRN5 chimeras bearing VIN3 domains were highly inefficient in coIP HA-PRC2 (**Fig. 3E**, lanes 1-8), while the gradual substitutions of domains in GFP-VIN3 with the corresponding VRN5 domains restored PRC2 coIP in the assays, as expected (**Fig. 3E**, lanes 9-15). This is particularly striking in the case of the FN3 substitution in GFP-VIN3 (**Fig. 3E**, lane 11), which suggests that FN3 may confer the VRN5-specific interaction with PRC2 (see also below). In addition, the linker of VRN5 also contributes to its association with MSI1 (**Fig. 3E**, lane 14) and, together with PHDsuper and FN3, imparts efficient association with the whole PRC2 (and particularly with its VRN2 scaffold; **Fig. 3E**, lane 2 vs 15). These results suggest that VRN5 not only binds to the MSI1 subunit of PRC2, but also uses its PHDsuper-FN3 module to contacts its VRN2 scaffold, whereby the latter may determine the binding preference of PRC2 for VRN5 (see below).

Finally, we note that the VRN2-PRC2 complex lacks the C2 domain present in SUZ12, the human counterpart of VRN2 (Gendall et al. 2001). Interestingly, this C2 domain contributes to the binding site of the mammalian PRC2 complex for its accessory factors PHF19 or AEBP2 (Chen et al. 2018; Chen et al. 2020). Strikingly, FN3 is a close structural relative of C2 (**Supplemental Fig. S5G,H**), suggesting that the plant FN3 domain in accessory protein VRN5 mimics the C2 domain in the core PRC2 subunit in the human complex. Based on the results from our mutational analysis (**Fig. 3E**) and bearing in mind the structure of the human PRC2 complex (Kasinath et al. 2018), we propose that the VRN5 FN3 domain interacts mainly with the regulatory N-lobe of the VRN2 scaffold and with the MSI1 subunit of the plant PRC2 complex (see below). Further structural studies are required to fully elucidate these functional parallels.

### Intramolecular interactions between VRN5 PHDsuper and FN3

Our attempts to determine the crystal structure of the VRN5 PHDsuper-FN3 module from Arabidopsis and other plant species (including palm tree and maize) following expression in *E.coli* were unsuccessful. We therefore resorted to cross-linking mass spectrometry (XL-MS) analysis to test whether we could detect an interaction between PHDsuper and FN3. We purified a PHDsuper-FN3 fragment from Zm VRN5 (because of the relatively high yields of the maize protein) and prepared the purified protein for XL-MS. This revealed multiple intra-molecular cross-links between FN3 and the hinge region, within FN3 itself and within the 4HB module of PHDsuper (**Supplemental Fig. S6A,B**). Repulsive point mutations in key residues mediating these cross-links (H297E and E299R) blocked several of these cross-links, leading to an increased degree of freedom between the constituent domains. This is even more pronounced if a third repulsive mutation is introduced, which caused large (>27Å) structural alterations. Finally, if four predicted interface residues were mutated (H297E, E299R, R216E, E260R), most of the inter-domain cross-links disappeared, whereby the only remaining ones were within the 4HB module of PHDsuper (**Supplemental Fig. S6C-E**). Overall, our XL-MS results support the notion of a substantial structural change following mutational disruption of the predicted interface between PHDsuper and FN3.

The deep conservation of the four key residues within VRN5 FN3 (designated key quartet) that mediate its intramolecular interaction with PHDsuper (**Fig. 4A; Supplemental Figs. S2, S6**) indicates that the resulting compact conformation is inherent to all angiosperm VRN5 orthologs. To test the functional relevance of these FN3 residues, we introduced repelling point mutations into Arabidopsis GFP-VRN5 and tested the association of these mutants with PRC2 by coIP (**Fig. 4B**). Remarkably, each of them strongly reduced the PRC2 coIP with GFP-VRN5 (**Fig. 4B**, lanes 2-8), similarly to deletion of the β14-strand of FN3 (Δβ14) (**Fig. 4B**, lane 9) that appears to engage in several close interactions with partner residues. Neither the key quartet nor β14 are conserved in VIN3 and VEL1 paralogs (**Supplemental Fig. S3**), consistent with our notion that the compact conformation conferred by these residues is unique to VRN5 orthologs. In summary, the compact conformation of Arabidopsis VRN5 described above is a defining structural feature of VRN5 orthologs that is functionally relevant for binding to PRC2.

**Figure 4.**
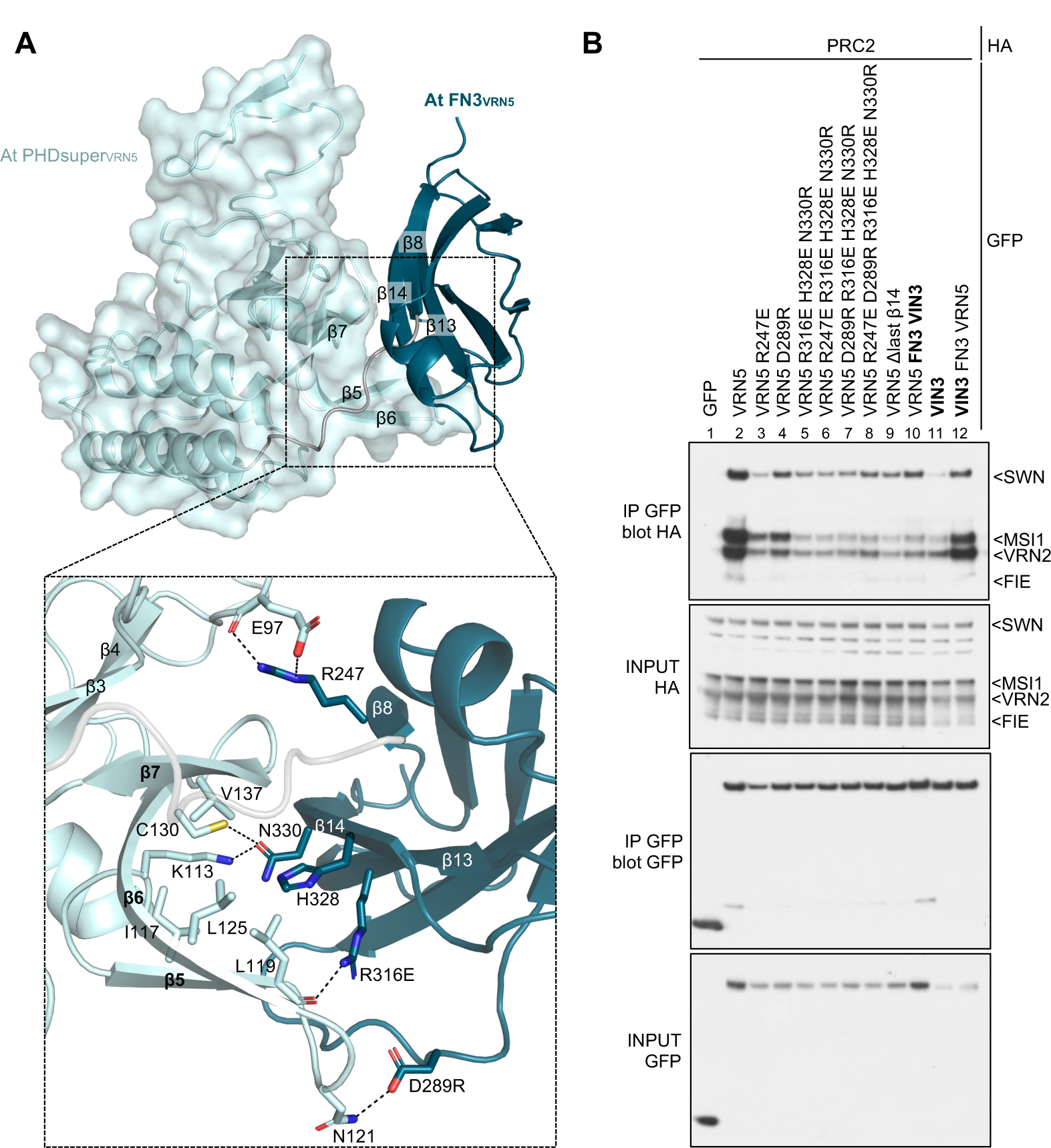
Analysis of intramolecular inter-domain interactions within VRN5. (A) Structural prediction of At VRN5 PHDsuper-FN3_VRN5_ (in ribbon and surface representations), with key structural elements mediating inter-domain interactions highlighted; *light blue*, PHDsuper; *teal*, FN3 domain. Close-up view of PHDsuper-FN3 interactions, with key interacting residues in *stick* representation; *dashed lines*, hydrogen bonds or salt bridges. (B) coIP of HA-tagged PRC2 core components with wt or selected inter-domain mutants of GFP-VRN5 (indicated above panel), as in Fig. 2A.

### VRN5 binds predominantly to the regulatory lobe of PRC2

The structure of the Arabidopsis PRC2 complex predicted by Alphafold2 (Jumper et al. 2021) consists of a catalytic module comprising the histone methyltransferase SWN, FIE and the C-terminus of VRN2 (C-lobe) plus a regulatory module containing MSI1 and the N-terminus of VRN2 (N-lobe) (Chen et al. 2018; Kasinath et al. 2018; Grau et al. 2021). To establish the importance of each PRC2 subunit for their interaction with VRN5, we co-expressed GFP-VRN5 with all but one of the four PRC2 subunits for coIP assays. This revealed that VRN2 and MSI1 are crucial for the coIP of PRC2 with GFP-VRN5 (**Fig. 5A**, lane 3 and 6) whereas FIE is not (**Fig. 5A**, lane 5). Moreover, SWN is the only PRC2 subunit that associates with GFP-VRN5 independently of the others (**Fig. 5A**, lane 8 versus 4). Next, in coIP assays of GFP-VRN5 co-expressed with combinations of two individual PRC2 modules, we found that VRN5 associates with the two components of the regulatory module, namely MSI1 and the N-lobe of VRN2 (**Fig. 5B**). This suggests an extensive interface of VRN5 with the regulatory PRC2 module.

**Figure 5.**
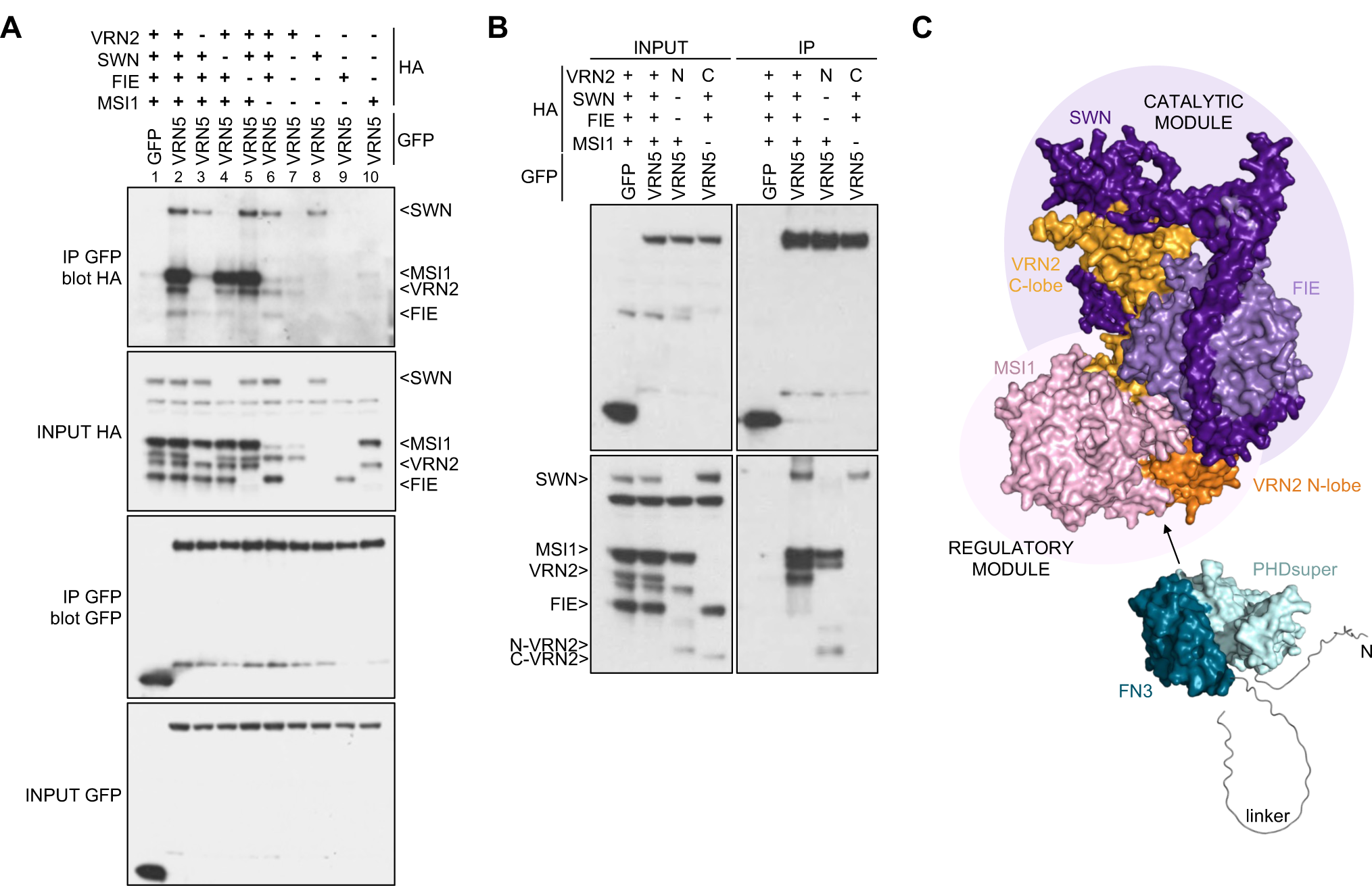
Interaction between VRN5 and lower module of PRC2. (A) coIP of different combinations of HA-tagged PRC2 with GFP-VRN5, as in Fig. 2A. (B) coIP of HA-tagged catalytic or regulatory modules of PRC2 with GFP-VRN5, as in Fig. 2A; *N*, VRN2 N-lobe; *C*, VRN2 C-lobe. (C) Model of interaction between structure prediction of VRN5 and At PRC2 complex, with catalytic module (*purple background*) comprising SWN (*purple*), FIE (*light purple*) and VRN2 C-lobe (*light orange*) and regulatory module (*light pink background*) comprising MSI1 (*pink*) and VRN2 N-lobe (*orange*).

To test this notion, we performed XL-MS analysis of the purified VRN5ΔVEL-PRC2 complex (**Supplemental Fig. S7**). This revealed multiple cross-links between MSI1 and the N-terminal region of VRN5 that precedes its PHDsuper, but also between the VRN5 4HB module and SWN and MSI1. Moreover, we found several cross-links between MSI1 and the upstream part of the unstructured linker region of VRN5, and between the latter and the N-terminal lobe of VRN2. Importantly, we found no cross-links between VRN5 and FIE, consistent with our coIP data (**Fig. 5A, B**). Taken together, our XL-MS and coIP results identify the regulatory module of PRC2 as the binding site for VRN5 (**Fig. 5C**), although further analysis will be required to determine the precise details of the molecular interaction between the two.

### VAL1 is dispensable for recruiting VIN3 to the FLC nucleation region

One boundary of the nucleation region, where VIN3 and VRN5 co-localize and where H3K27me3 accumulates, is the binding site for VAL1. VAL1 is a transcriptional repressor that acts as an assembly platform for co-transcriptional repressors and chromatin regulators (Questa et al. 2016). VAL1 has a PHD-L domain, a DNA sequence-specific binding domain (B3), a CW domain and a ZnF-EAR motif (**Fig. 6A**). Given the direct physical link between VAL1 and *FLC*, we asked whether we could detect a physical interaction between VAL1 and any of the VEL paralogs in coIP assays. Indeed, monomeric dsRed-VIN3 showed robust coIP with co-expressed GFP-VAL1, whereas dsRed-VRN5 and dsRed-VEL1 showed very little (**Fig. 6B**). This VIN3-VAL1 association depends on the linker region and CW domain of VAL1 (**Fig. 6C,D**), whereby the CW domains is a PHD-like zinc finger whose human homologs bind to H3K4me (Liu et al. 2016). It also depends on a short KRFK motif within VIN3 within the flexible linker separating FN3 and VEL, and additional flanking sequences in both proteins also contribute to a robust VIN3-VAL1 interaction (**Fig. 6E; Supplemental Fig. S8A-E**).

**Figure 6.**
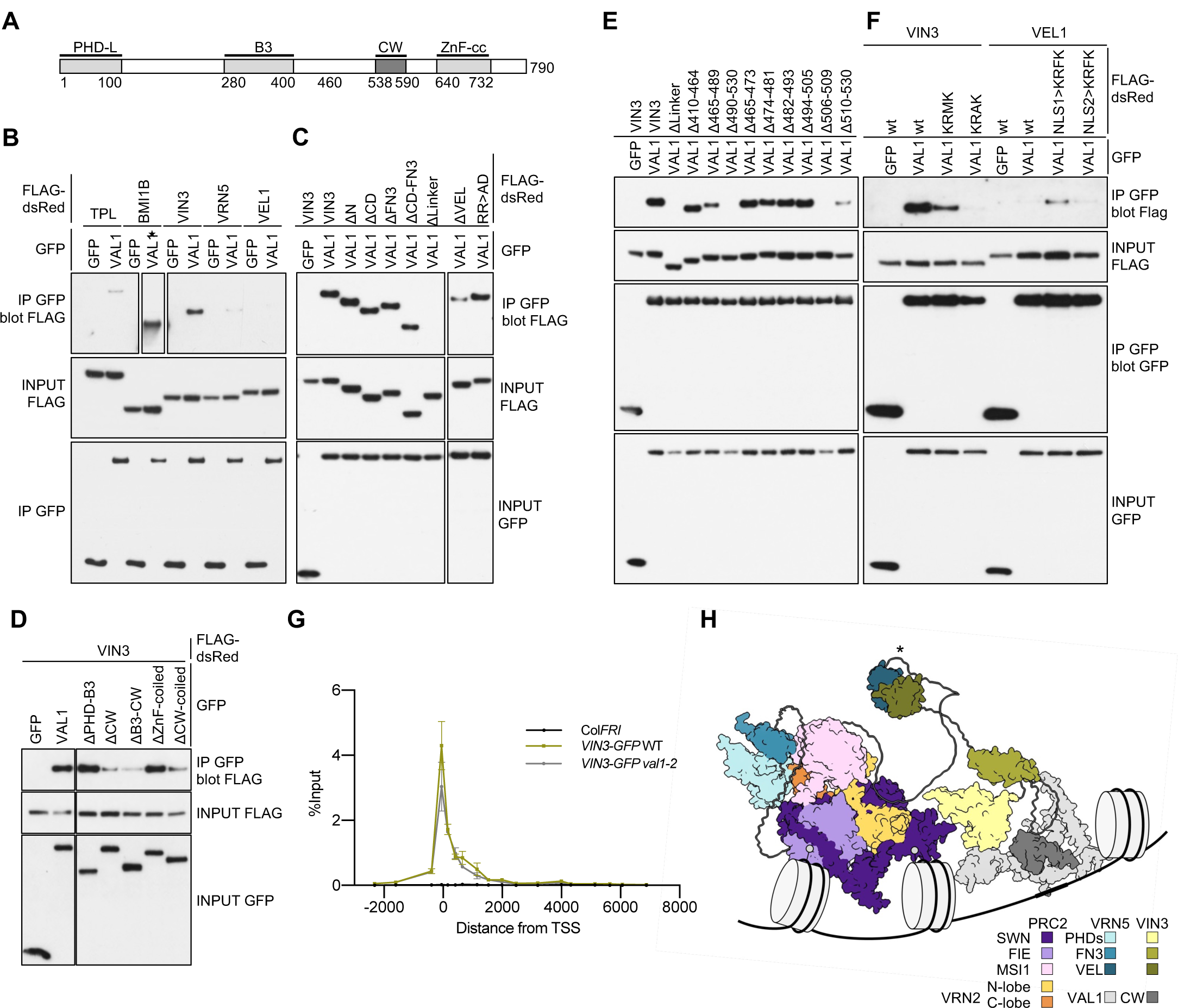
Association between VIN3 and VAL1. (A) Domain architecture of VAL1; *dark grey*, VIN3-interacting CW finger. (B-F) coIP of GFP-tagged wt or internal deletions of VAL1 with positive controls TPL and BMI1B FLAG-dsRed-tagged (star indicates short exposure) and wt or mutant FLAG-dsRed-tagged VEL proteins, as indicated above panels, as in Fig. 2A, revealing the critical role of VIN3 KRFK (Δ506-509) in (E); note also that KRFK suffices to confer some VAL1 interaction on VEL1 (F), see also text. (G) VIN3 enrichment at *FLC* in wt and *val1-2* mutant Arabidopsis vernalized for 6 weeks (6WT0), with non-transgenic *Col-FRI* as a negative control. Data are shown as percentage input; *error bars*, means ± s.e.m. from two independent experiments. (H) Model based on AlphaFold2 predictions of the VRN5-PRC2 complex and its interaction with VIN3-VAL1 mediated by a heterotypic VEL-VEL interaction (asterisk), with VAL1 being bound to the nucleation region of *FLC*; for simplicity, some flexible regions are not shown.

Intriguingly, the KRFK motif conforms to a classical monopartite nuclear localization signal (cNLS) (K K/R X K/R) (Chelsky et al. 1989), which confers strong and specific binding between cNLS-containing cargo and importin (Conti and Kuriyan 2000; Hodel et al. 2001; Marfori et al. 2012). Consistent with this, deletion of KRFK renders dsRed-VIN3 cytoplasmic but its nuclear localization and association with VAL1 can be restored by a heterologous viral NLS (PKKKRKV) inserted next to the N-terminal tag (**Supplemental Figs. S8, S9**), perhaps because of the sequence resemblance between the two motifs. Importantly however, substitution of KRFK in dsRed-VIN3 with KRMK or KRAK strongly reduces, or abolishes, its coIP with GFP-VAL1 (**Fig. 6F**), although both substitutions retain their nuclear localization (**Supplemental Fig. S9**), as expected since the third is the least important residue in cNLS motifs as regards their affinities to importin (Hodel et al. 2001; Marfori et al. 2012). By contrast, the third residue within KRFK is clearly a critical determinant for the association of VIN3 with VAL1.

VEL1 contains two putative cNLS motifs, one upstream of its PHD finger (KRQR) and another downstream of it (KRMK; **Supplemental Fig. S8F**), yet VEL1 poorly coIPs with GFP-VAL1 (**Fig. 6B**). Again, this argues against the notion that the nuclear localization of VIN3 *per se* enables it to bind to VAL1. In further support of this, dsRed-VEL1 mutants bearing a KRFK substitution of either motif alone does not affect their nuclear localization (**Supplemental Fig. S9**), but the upstream albeit not the downstream KRFK substitution confers weak yet readily detectable coIP with GFP-VAL1 (**Fig. 6F**). Evidently, KRFK is sufficient to confer modest VAL1 association on VEL1, albeit in a context-sensitive manner (note that the downstream substitution may be too close to FN3 to allow it to interact with VAL1) (**Supplemental Fig. S8F**).

VIN3-GFP is associated with the nucleation region in control plants vernalized for 6 weeks (6WT0) (Yang et al. 2017). We therefore tested whether this was reduced in *val1-2*. Interestingly, loss of VAL1 did not change VIN3 association at the nucleation region (**Fig. 6G**). Thus, *in vivo* association of these two proteins may be highly dynamic or part of a multifactorial assembly complex, where other factors can substitute for loss of VAL1. The observation that VIN3 is dispensable for the deposition of H3K27me2 at *FLC* (**Fig. 1B**) and that VAL1 is dispensable for VIN3 tethering at *FLC* (**Fig. 6G**) imply that the methyl transferase activity of VRN2-PRC2 can be targeted to the *FLC* nucleation region in the absence of VAL1 or VIN3. In support of this notion, a mutation preventing binding of VAL1 (C585T) (Questa et al. 2016) still accumulates H3K27me2 at the *FLC* nucleation region (**Supplemental Fig. S10**). Presumably, this targeting event is achieved by a combination of factors that bind directly to this key cis-regulatory region of *FLC*. Thus, this region can be likened to the Polycomb response elements in Drosophila that are cis-linked to PRC2 target genes and known to contain binding sites for multiple DNA-binding proteins (Kassis and Brown 2013).

Given that VIN3-dependent polymerization is required for the silencing of *FLC* in the cold (Fiedler et al. 2022) and our new results suggesting that VIN3 facilitates the PRC2-mediated conversion of H3K27me2 to H3K27me3, a potential model is that VIN3 facilitates stable tethering of VRN5-PRC2 to *FLC* to facilitate the slow, gradual conversion of H3K27me2 to H3K27me3, possibly through VEL-dependent co-polymerization with VRN5 (**Fig. 6H**).

### VEL proteins co-localize with H3K27me3 at numerous sites throughout the Arabidopsis genome

To pursue the generality of VEL protein function and how they connect with PRC2 silencing we undertook ChIP-seq analysis using lines expressing VIN3-GFP, VRN5-YFP, or VEL1-3xFLAG, in either non-vernalized plants (NV) or after 6 weeks of cold (6WT0). We only considered highly significant peaks (q-value ≤ 10^!’#^) in our analysis. This identified many more potential target genes for VEL1 than for VIN3 and VRN5: 8357 and 7647 genes enrich for VEL1 binding at NV and 6WT0 respectively. We identified 49 (NV) and 1242 (6WT0) potential target genes for VIN3, and 1130 (NV) and 2728 (6WT0) for VRN5 (**Fig. 7A**). VEL1 may thus have a broader role in the Arabidopsis genome, or the different tags (FLAG on VEL1 versus GFP or YFP on VIN3 or VRN5, respectively) contribute to the different enrichments of these proteins. We favor the former explanation as visual inspection revealed that the vast majority of “VEL1 only peaks” also show lower than threshold VIN3 and VRN5 enrichment. The genome-wide trend is that all three VEL proteins co-occupy the same genomic region (**Fig. 7B**). After 6 weeks of cold, an additional 1193 and 1598 significant potential target genes were identified for VIN3 and VRN5, respectively (**Fig. 7A**). Increased VIN3 levels thus increase the residency time of not only VIN3 but also VRN5 at many target genes, including *FLC* (**Supplemental Fig. S11A**). Consistent with this, an additional 1497 genes have recently been reported to have increased H3K27me3 levels after cold exposure (Xi et al. 2020). One example is *RSH3* (At1G54130), which has been linked to downregulation of chloroplast transcription in response to abscisic acid (Yamburenko et al. 2015). Like *FLC*, *RSH3* shows an obvious increase, particularly of VIN3 but also of VEL1 and VRN5 (**Fig. 7C**).

**Figure 7.**
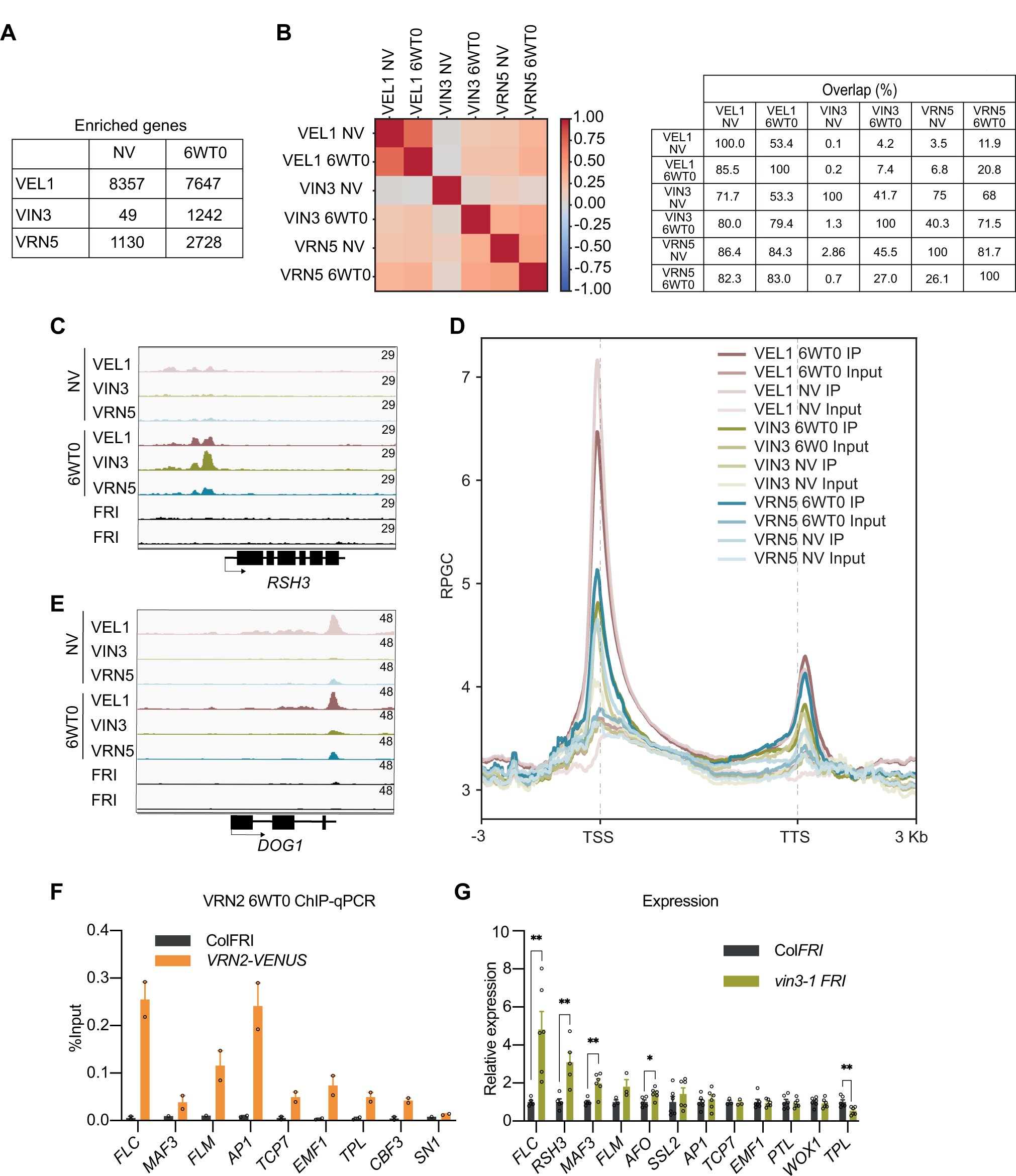
Genome-wide occupancy of VEL1, VIN3, and VRN5. (A) Table showing the number of VEL1, VIN3, and VRN5 target genes at NV and 6WT0. Target genes present in at least two replicates were counted. (B) Heatmap showing the multiple overlaps among the VEL1, VIN3, and VRN5 at NV and 6WT0 peaks (left), and the percentage overlap between peaks identified for VEL1, VIN3, and VRN5 at NV and 6WT0 (right). (C) IGV screenshots showing the co-localization of the VEL proteins at the TSS of *RSH3*. (D) Metagene plots of VELs distribution over transcription units and flanking regions. TSS, transcription start site; TTS, transcription termination site. RPGC, reads per genomic content. Inputs from the respective VEL IPs are used as controls. (E) IGV screenshots showing the co-localization of the VEL proteins at the TTS of *DOG1*. (F) ChIP-qPCR showing enrichment of VRN2 after six weeks of cold treatment (6WT0) at ten potential target genes. Data are shown as the percentage input. Non-transgenic ColFRI plants were used as a negative control sample. *FLC* was used as a positive control locus, and *SN1* was used as a negative control locus. Error bars are means ± s.e.m. from two independent experiments. (G) Expression of several potential target genes in the mutant *vin3-1 FRI* relative to the wild-type Col*FRI* at 6WT0. Data are shown normalized to the expression level of the respective gene in Col*FRI*. Error bars are means ± s.e.m. from at least three biological replicates. For statistical tests, a single asterisk denotes p<0.05, and two asterisks denote p<0.01 between samples by Student’s t-test.

The VEL proteins preferentially enrich at transcription start sites (TSS) (**Fig. 7D**) with a smaller but significant peak frequently detected slightly downstream of the transcription termination site (TTS) (**Fig. 7D**). For example, at *DELAY OF GERMINATION 1* (*DOG1*), there is strong enrichment of VEL1, and to some extent of VIN3 and VRN5 at the TTS (**Fig. 7E)**. The VEL genome-wide pattern is similar to the core PRC2 proteins SWN and CLF (Shu et al. 2019), consistent with VEL proteins acting as PRC2 accessory factors.

Comparison of the VEL ChIP-seq data with published genome-wide Arabidopsis H3K27me3 datasets, focusing on the 869 potential VEL target genes common between VEL1 and VRN5 in NV conditions, showed around 23% overlap: 162 (Zhang et al. 2007), 180 (Shu et al. 2019), 207 (Li et al. 2015) or 213 (Zhou et al. 2017) genes are common between H3K27me3 marked genes and genes bound by VEL1 and VRN5 (**Supplemental Fig. S11B**). Interestingly, almost all of these target genes overlap with the PRC1-mediated modification H2AK121ub (**Supplemental Fig. S11C**) (Zhou et al. 2017). 570 of the 869 VEL1 or VRN5 target genes overlap with genes marked by H2AK121ub (**Supplemental Fig. S11C**), suggesting a tight link between PRC1-mediated H2AK121ub and chromatin occupancy of VEL proteins.

The functional significance of the ChIP-seq data is best elaborated at the well-established PRC2 target *FLC* and its *MAF1-MAF5* relatives (Ratcliffe et al. 2001; Ratcliffe et al. 2003). At *FLC*, VEL1, VIN3 and VRN5 are found in the nucleation region as expected for a 6WT0 treatment (**Supplemental Fig. S11A**) (Yang *et al*, 2017). The enrichment of VEL1 resembles the pattern for VRN5, enriched at the nucleation region in NV conditions, and becoming more enriched with cold exposure (**Supplemental Fig. S11D**). *MAF1/FLM, MAF2* and *MAF3* are repressed by vernalization, while *MAF4* and *MAF5* are not (Sheldon et al. 2009; Kim and Sung 2013); *MAF5* is induced by vernalization (Ratcliffe et al. 2003). Consistent with this, we found VIN3 was enriched at *FLM*, *MAF3*, and to some extent at *MAF2* (**Supplemental Fig. S11A**), but not at *MAF4* and *MAF5*. VRN5 was enriched at *FLM*, but not at *MAF2*-*MAF5*, consistent with *vrn5* mutants mostly affecting *FLC* expression and to some extent *FLM*, but not other members of the *FLC* gene family (Kim and Sung 2013). Likewise, VEL1 is enriched at *MAF4* and *MAF5*, in addition to *FLC* and to some extent *FLM* (**Supplemental Fig. S11A**), consistent with *MAF4* and *MAF5* being reported to be mis-regulated in *vel1* mutants (Kim and Sung 2013). We note that some enrichment is also found at the 3’ end of *MAF5*, which might reflect the different behaviour of *MAF5* during vernalization compared to the other *FLC* clade members. These differential enrichments point to different importance of VEL proteins in PRC2 silencing at different loci.

We used ChIP-qPCR to validate an additional selection of the targets identified by ChIP-seq (**Supplemental Fig. S11E**). In addition, to link their regulation to PRC2, we also tested for co-localization of VRN2 using a transgenic line expressing VENUS-tagged VRN2 (VRN2-VENUS) that complements the *vrn2-1* mutant. VRN2-VENUS enrichment was detected at all selected VEL targets (**Fig. 7F**). Genes showing strong enrichment of VIN3 were also frequently mis-regulated in a *vin3* mutant background (**Fig. 7G**), and to a lesser extent in the *vrn5* mutant (**Supplemental Fig. S11F**). Consistent with VIN3 working with PRC2, in four out of five cases we observed a release of repression in the *vin3-1* mutant, similar to previous studies analysing mis-regulation in PRC2-defective mutants (Shu et al. 2019).

### Conclusions

Here, we find that different VEL proteins (cold-induced VIN3, constitutively expressed VRN5 and VEL1) constitute an evolutionarily conserved set of plant proteins that have functionally distinct activities important for Polycomb silencing. VRN5 directly and extensively interacts with multiple subunits of PRC2, enabling H3K27me2 and H3K27me3 deposition at the nucleation region of *FLC*. VEL1 may contribute to this role. By contrast, VIN3 interacts with VAL1, as part of a multifactorial assembly platform coordinating various activities for epigenetic silencing of *FLC*, likely by connecting PRC1 and PRC2 activities (Mikulski et al. 2022) and facilitating the PRC2-mediated conversion of H3K27me2 to H3K27me3. Homo-and potentially heteropolymerization of VIN3, VRN5 and VEL1 VEL domains could promote dynamic biomolecular condensation of Polycomb complexes (Fiedler et al. 2022), thereby imparting a high binding avidity on these complexes (Bienz 2014; Pancsa et al. 2019; Bienz 2020). Our model is that VEL proteins work collectively, contributing to PRC2 stabilization and association, enabling the transition from H3K27me2 to H3K27me3. Although this mechanism has been elaborated predominantly at the *FLC* locus, the evolutionary conservation of proteins and the genome-wide co-localization of the VEL proteins with H3K27me3 targets suggests a broad role for VEL proteins in plant growth and development.

## Material and methods

### Generation of plasmids

VEL sequences (VIN3, Q9FIE3; VEL1, Q9SUM4; VRN5, Q9LHF5) for in vitro and cell-based assays were amplified by polymerase chain reaction (PCR) from either plasmid templates (Greb et al. 2007) synthetic genes (gBlocks, IDT), cloned into mammalian or bacterial expression vectors by restriction-free cloning. Point mutations and deletions were generated by Quikchange, using KOD DNA polymerase (Merck Millipore). All plasmids were verified by sequencing.

### Protein expression and purification

PHDsuper-FN3_VRN5_ *from* Zm for XL-MS analysis was inserted into pEC-LIC-His-3C containing a hexa-histidine tag and 3C protease cleavage site at the N terminus and was expressed in BL21 CodonPlus (DE3)-RIL cells (Agilent) in LB medium. Cells were grown at 37°C to OD_600_ 0.6, then moved to 18°C, followed by induction with 0.4 mM IPTG and 100 μM ZnCl2 at OD_600_ 0.8. Harvested cells were resuspended in lysis buffer (25 mM Tris pH 8.0, 200 mM NaCl, 20 mM imidazole pH 8, and EDTA-free protease inhibitor cocktail (Roche) and lysed by sonication (Branson). Cleared lysates were loaded onto Ni-NTA resin (Qiagen) and washed with lysis buffer. After extensive washing, samples were eluted with lysis buffer supplemented with 300 mM imidazole. Eluted samples were incubated with 2 mM DTT and cleaved by 3C protease (made in house; protease:protein ratio 1:80) overnight at 4°C. After overnight incubation with 3C, sample was loaded onto a HiLoad 26/600 Superdex 200 pg column (GE Healthcare) equilibrated in 25 mM Tris pH 8, 150 mM NaCl, and 1 mM DTT. All the purifications process were done at 4°C.

Pd PHD_VIN3_ patch mutants (patch 1: residues R142E, R154E, K167E and K233E, patch 2: residues R242E, R243E, R294E and R299E, and patch 3: residues K197E, K256E, K265E and K276E) were expressed and purified as described before (Franco-Echevarría et al. 2022).

At VRN5Δ1VEL-PRC2 complex was co-expressed in SF9 insect cells using biGBac method (Weissmann et al. 2016). N-terminal strep-tag II tagged VRN5ΔVEL (minimal VRN5 construct that showed the strongest PRC2 interaction in our coIP experiments), N-terminal HA-tagged VRN2_1-397_, SWN_29-831_ and FIE_1-360_, and C-terminus HA-tagged MSI1_1-412_ were cloned into a pLIB vector, separately. The five genes were subsequently sub-cloned into a pBIG1a vector by a Gibson assembly reaction, in which these gene PCR products were connected in series with the linearized pBIG1a vector digested by SwaI. The recombinant baculovirus was generated using FuGENE HD Transfection reagent (Promega) in Sf9 cells using Insect X-press (Lonza) medium and infected cells were grown for 60-72h at 27 °C before harvesting them for protein extraction and purification. Harvested cells were resuspended in 25mM HEPES pH8, 250mM NaCl, 2mM MgCl_2_, 1mM DTT, 5% glycerol and EDTA-free protease inhibitor cocktail (Roche) and lysed by sonication. After centrifugation at 15,000 rpm for 30 min, the supernatant was incubated with Strep-tactin superflow resin (Qiagen) for 2 hours and then extensively washed with lysis buffer and eluted in lysis buffer with 5mM desthiobiotin and further purified by size-exclusion chromatography with Superose 6 increase 10/300 GL-column (GE Healthcare).

### Phylogenetic analysis

Protein sequences of VEL orthologs were obtained from BLAST (Camacho et al. 2009) or retrieved from JACKHMMER (https://www.ebi.ac.uk/Tools/hmmer/search/jackhmmer). Alignments of sequences were done with Mac-Vector (MacVector Inc) and ESPRIPT using Clustal Omega (https://www.ebi.ac.uk/Tools/msa/clustalo/).

### Alphafold2 predictions

Structure predictions and PAE plots were calculated from ColabFold AlphaFold2 using MMseqs2 (https://colab.research.google.com/github/sokrypton/ColabFold/blob/main/AlphaFold2.ipynb).

### XL-MS

Protein cross linking reactions were carried out at room temperature for 60 minutes with 50 mg of complex present in 1 mM, 5 mM, 10 mM, and 20 mM of EDC. Crosslinked protein was quenched with the addition of Tris buffer to a final concentration of 50 mM. The quenched solution was reduced with 5 mM DTT and alkylated with 20 mM idoacetamide. SP3 protocol as described in (Hughes et al. 2019) (Batth et al. 2019) was used to clean-up and buffer exchange the reduced and alkylated protein, shortly; proteins are washed with ethanol using magnetic beads for protein capture and binding. The proteins were resuspended in 100 mM NH_4_HCO3 and were digested with trypsin (Promega, UK) at an enzyme-to-substrate ratio of 1:25, and protease max 0.1% (Promega, UK). Digestion was carried out overnight at 37°C. Clean-up of peptide digests was carried out with HyperSep SpinTip P-20 (ThermoScientific, USA) C18 columns, using 80% Acetonitrile as the elution solvent. Peptides were then evaporated to dryness via Speed Vac. Samples were fractionated via size exclusion chromatography using a Superdex 30 Increase 3.2/300 column (GE heathcare, Sweden) at a flow rate of 20 uL/min of 30% ACN 0.1 % TFA. Fractions are taken every 3 minutes, and the fractions 1-3, 4-6, were concatenated and fractions 5 to 10 were analyzed separately, all samples were dried on speed-vac. Dried peptides were suspended in 3% Acetonitrile and 0.1 % formic acid and analysed by nano-scale capillary LC-MS/MS using an Ultimate U3000 HPLC (ThermoScientific, USA) to deliver a flow of 300 nl/min. Peptides were trapped on a C18 Acclaim PepMap100 5 μm, 100 μm x 20 mm nanoViper (ThermoScientific, USA) before separation on PepMap RSLC C18, 2 μm, 100 A, 75 μm x 50 cm EasySpray column (ThermoScientific, USA).

Peptides were eluted on a 110 minute gradient with acetonitrile and interfaced via an EasySpray ionisation source to a quadrupole Orbitrap mass spectrometer (Q-Exactive HFX, ThermoScientific, USA). MS data were acquired in data dependent mode with a Top-10 method, high resolution scans full mass scans were carried out ((R = 120,000, m/z 400 – 1550) followed by higher energy collision dissociation (HCD) with stepped collision energy range 26, 30, 34 % normalised collision energy. The tandem mass spectra were recorded (R=60,000, AGC target = 1 x 105, maximum IT = 120 ms, isolation window m/z 1.6, dynamic exclusion 50 s). Cross linking data analysis: Xcalibur raw files were converted to MGF files using ProteoWizard (Chambers et al. 2012) and cross links were analysed by MeroX (Gotze et al. 2015). Searches were performed against a database containing known proteins within the complex to minimise analysis time with a decoy data base based on peptide sequence shuffling/reversing. Search conditions used 3 maximum missed cleavages with a minimum peptide length of 5, cross linking targeted residues were K, S, T, and Y, cross linking modification masses were 54.01056 Da and 85.98264 Da. Variable modifications were carbmidomethylation of cysteine (57.02146 Da) and Methionine oxidation (15.99491 Da). False discovery rate was set to 1 %, and assigned cross linked spectra were manually inspected.

### coIP assays in human cells

Protein co-immunoprecipitation assays were carried out in HEK293T cells grown on DMEM supplemented with 10% FBS, seeded on poly-L-lysine coated plates at ∼70% confluency, 89 mm tissue culture dishes per coIP was used. After attachment, cells were transfected with a DNA:PEI (1:3.5) mixture. With the purpose of achieving near stoichiometric expression of each protein in PRC2-VRN5 coIPs, 10 μg of total DNA was used per culture dish (2 μg of GFP-VRN5, 4.3 μg of VRN2, 2.4 μg of SWN, 0.3 μg of MSI1 and 1 μg of FIE). For VAL1 and VEL proteins coIPs, 8 μg total DNA (2 μg of GFP-VAL1 and 6 μg of FLAG-dsRed-TPL/BMI1B (dsRed-monomer N1 vector), or VEL proteins). Cells were lysed ∼18 h post-transfection in lysis buffer (20 mM Tris pH 7.4, 200 mM NaCl, 10% glycerol, 5 mM NaF, 2 mM Na3VO4, 1mM EDTA, 0.2% Triton X-100, EDTA-free protease inhibitor cocktail (Roche)). Lysates were clarified by centrifugation (16100 rcf, 20 min), and supernatants were incubated with GFP-trap (Chromotek) for 90 min at 4°C on an over-head tumbler. Immunoprecipitates were washed three times in lysis buffer and eluted by boiling in LDS sample buffer for 10 min. Input and coIP fractions were separated by SDS-PAGE, blotted onto PVDF membrane, checked for equal loading by Ponceau staining, and processed for Western blotting with appropriate antibodies. Primary antibodies (anti-GFP (Sigma, #G1544), anti-HA (Abcam, #ab9110) or anti-FLAG (Sigma, #7425)) and secondary antibodies were diluted 1:5000 in PBS, 0.1% Triton X-100 and 5% milk powder. Blots were washed with PBS, 0.1% Triton X-100 and developed with ECL Western Blotting Detection Reagent on film.

### Immunofluorescence

HEK293T cells were transfected with 1 mg total DNA and 3.5x PEI. HEK293T cells were transfected with 1 μg total DNA (250 ng of GFP-VAL1 and 750 ng of FLAG-dsRed-VIN3 and FLAG-dsRed-VEL1 constructs) and expressed for 18 hours. PBS washed cells were fixed on coverslips with 4% formaldehyde in PBS for 20 minutes and subsequently permeabilised by 0.5% Triton-X100 in PBS for two minutes. Coverslips were washed with PBS-T and embedded with VectaShield with DAPI mounting media. Images were acquired with identical settings using a Zeiss 710 Confocal Microscope using ‘best signal’ setting (Smart Setup, ZEN software, Zeiss).

### Plant materials and transgenic constructs

The transgenic lines pVIN3:VIN3-eGFP/*vin3-4* (VIN3-GFP), pVRN5:VRN5-YFP/*vrn5-8* (VRN5-YFP), pFLC:FLC-WT/*flc-2* FRI and pFLC:FLC-C585T/*flc-2* FRI were described previously (Greb et al. 2007; Questa et al. 2016; Yang et al. 2017). The VIN3-GFP *vin3 vrn5* line was obtained by crossing pVIN3:VIN3-GFP *vin3-4* FRI^SF2^ into the vrn5-8 FRI^SF2^ mutant (SALK_136506, crossed into Col-FRI^SF2^). Primers used for PCR genotyping to identify homozygous plants are listed in Table S1. The VEL1-3xFLAG construct was generated for this study by classic PCR cloning. The construct encodes *VEL1* genomic DNA from −2100 bp upstream of the ATG to 720 bp downstream of the stop codon. The 3xFLAG tag was inserted at the C-terminus by PCR. The construct was cloned into the binary vector pCAMBIA1300 by restriction cloning and transformed into *vel1-1* FRI^SF2^/fri. After identification of plants that contain the transgene, plants were made homozygous for FRI^SF2^ to give rise to VEL1-3xFLAG/*vel1* FRI^SF2^. To generate the VRN2-Venus line, the genomic *VRN2* sequence from Col-0 was cloned from −1364 up upstream of the ATG and fused with C-terminal Venus excluding the native VRN2 stop codon. T3A was used as a terminator sequence. The construct was cloned into the binary vector pCAMBIA1300 by restriction cloning and transformed into *vrn2-1* Col-FRI (Yang et al. 2017) mediated by Agrobacterium C58 using the floral dip method.

### coIP assays in plants

Total proteins were extracted from 6 g of frozen ground Arabidopsis seedling tissue with IP buffer (50 mM Tris-HCl pH 7.5, 150 mM NaCl, 0.5% NP-40, 1% Triton, EDTA-free protease inhibitor cocktail [Roche]). Lysates were cleared by centrifugation (6,000 g, 30 min, 4°C) and incubated with GFP-Trap (Chromotek) for 2h. Immunoprecipitates were washed four times with IP buffer and eluted by boiling in 4x NuPAGE LDS sample buffer for 10 min. Input and coIP fractions were separated by polyacrylamide gel electrophoresis (SDS-PAGE) and blotted onto polyvinylidine difluoride (PVDF) membranes. Primary antibodies anti-GFP (11814460001, Roche) and anti-FIE (AS12 2616, Agrisera) were diluted 1:1000, anti-Actin was diluted 1:5000 (AS132640, Agrisera). Secondary antibodies were HRP-coupled. Blots were washed with TBS containing 0.05% Tween-20 and developed with SuperSignal West Femto Maximum Sensitivity Substrate (Thermo Scientific).

### ChIP and ChIP-seq

For all ChIP analyses, plant material was cross-linked by vacuum-infiltrating seedlings in a solution of 1% (w/v) formaldehyde in PBS buffer for 10 min. The cross-linking reaction was stopped by adding glycine to a final concentration of 0.125 M followed by 5 min of vacuum infiltration. Nuclei were extracted from 2 g of ground frozen material with Honda buffer (0.44 M Sucrose, 1.25% Ficoll, 2.5% Dextran T40, 20 mM Hepes KOH pH 7.4, 10 mM MgCl_2_, 0.5% Triton X-100, 5 mM DTT and Protease Inhibitors) and washed with Honda buffer several times. For histone ChIP, chromatin was extracted with nuclei lysis buffer (50 mM Tris/HCl pH 8, 10 mM EDTA, 1% [w/v] SDS, PIC) and sonicated using the Bioruptor Standard (Diagenode) for 15 cycles (30s on/ 30s off) to achieve an average fragment size of 200-500 bp. After removing cellular debris by centrifugation (10 min at 10.000 x *g*), the chromatin was diluted 10-fold with ChIP dilution buffer (1.1% [v/v] Triton x-100, 1.2 mM EDTA, 16.7 mM Tris/HCl pH 8, 167 mM NaCl). An aliquot corresponding to 1% (v/v) of the starting chromatin volume was removed as the input DNA control. Immunoprecipitation was performed with Dynabeads Protein A (Invitrogen) and α-H3K27me2 (Upstate, 07-452) or α-H3 antibody (abcam, ab1791) for 4h at 4°C on a rotator. Immunocomplexes were washed twice each with low salt wash buffer (150 mM NaCl, 0.1% [w/v] SDS, 1% [v/v] Triton X-100, 2 mM EDTA, 20 mM Tris/HCl pH 8), high salt wash buffer (500 mM NaCl, 0.1% [w/v] SDS, 1% [v/v] Triton X-100, 2 mM EDTA, 20 mM Tris/HCl pH 8) and TE buffer (1 mM EDTA, 10 mM Tris/HCl pH 8), and eluted twice with freshly prepared elution buffer (1% [w/v] SDS, 0.1 M NaHCO_3_) by incubating at 65°C for 15 min at 1000 rpm in a ThermoMixer. NaCl was added to the eluates and the input DNA aliquots to a final concentration of 0.2 M; samples were then incubated overnight (65°C, 600 rpm) for de-crosslinking and treated with proteinase K at 45°C for 1 h. DNA was purified with phenol/chloroform extraction and eluted into water. Histone ChIP samples were tested for enrichment by qPCR, with primer sequences for *FLC* and *Actin* (negative control) as previously published (Yang et al. 2017).

Non-histone ChIP followed by qPCR analysis was performed as described above with the following modifications: 3 g of plant material was cross-linked for 15 mins. After nuclei were resuspended in RIPA buffer (1x PBS, 1% IGEPAL CA-630, 0.5% Sodium deoxycholate, 0.1 % SDS, Roche Complete tables) and the DNA was fragmented to 200-500 bp by sonication. Anti-GFP (Abcam, ab290) and Protein A Agarose/Salmon Sperm DNA (Millipore, 16-157) were used for IP of GFP- and YFP-tagged proteins. For FLAG IP anti-FLAG (Sigma, F1804) was coupled to Dynabeads M-270 Epoxy (Invitrogen, 14301) following the manufacturer’s protocol (2 µl Anti-FLAG coupled to 1.5 mg Epoxy beads were used for each IP reaction). VRN2 ChIP was performed with the following further modifications: after a brief sonication to rupture nuclei, chromatin was treated with benzonase (0.002 U/µl chromatin; 70746-4 Millipore) at 4°C for 15 min to achieve a fragmentation of 200-500 bp. Benzonase was inactivated by adding EDTA to a final concentration of 10 mM before proceeding to IP. After IP, the EDTA concentration in all wash buffers was doubled.

Non-histone ChIP-seq was performed as described above with the modifications that rProtein A Sepharose Fast Flow (Merck, GE17-1279-01) was used instead of Protein A Agarose/Salmon Sperm DNA to avoid Salmon sperm DNA contamination. Each biological replicated was generated by pooling three individual ChIP pull-downs (each consisting of 6 g of whole seedling tissues) during DNA purification using the ChIP DNA Clean & Concentrator Kit (Zymo Research, D5201). The samples were sent to BGI Genomics for library preparation and sequencing using DNBSEQ. At least two biological replicates were sequenced for each genotype and conditions (n=3 for VIN3-eGFP 6WT0/VRN5-YFP NV/VRN5-YFP 6WT0/VEL1-3xFLAG NV/VEL1-3xFLAG 6WT0. n=2 for VIN3-eGFP NV/ColFRI-GFP IP NV/ ColFRI-GFP IP 6WT0/ ColFRI-FLAG IP NV and n=1 for ColFRI-FLAGIP 6WT0).

### ChIP-seq analysis

ChIP-seq reads were mapped to the TAIR 10 genome using the BWA (version 0.5.7) aligner with the parameters “-t 8 −l 25 −k 2 −n 5” and the resulting SAM alignment records were converted to BAM using the SAMtools (version 1.6). The peaks were called from the BAM output using the callpeak module from the MACS3 (version 3.0.0a7) (https://github.com/macs3-project/MACS) with the following parameters: “-q 0.05 −bdg −nomodel --extsize 180”. The peak and the corresponding BAM output was visualised using the Integrative Genome Viewer (IGV). The peaks overlapping the genes were identified using bedtools (version 2.27.1). The peaks were further processed and compared using the bespoke Python scripts are available at (https://github.com/threadmapper/VEL-ChIPseq).

## Data availability

The ChIP-seq raw data generated in this study has been deposited in the Sequence Read Archive (SRA) under the project: PRJNA973989.

## Competing interest statement

The authors declare no competing interests.

## Acknowledgements

We are indebted to Dr Danling Zhu for making the VEL1-FLAG transgene and Hongchun Yang for making the VRN2-VENUS transgene and homozygous transgenic line. The authors would also thank Shuqin Chen for their excellent technical assistance and Patricia Garay for thoughtful inputs for the ChIP-seq analysis. This work was funded by the European Research Council Advanced Grant (EPISWITCH, 833254), Welcome Trust (210654/Z/18/Z), BBSRC Institute Strategic Programmes (BB/J004588/1 and BB/P013511/1), and the Royal Society Professorship (RP\R1\180002) to C. D. and by the Medical Research Council (U105192713) to M. B.

## Author contributions

Conceptualization, M.B. and C.D.; investigation and data analysis, E.F.-E., M.N., A.S., J.C and T.E.M; writing – review & editing, all authors; supervision, M.B. and C.D.; funding acquisition, M.B. and C.D.; project administration, M.B. and C.D.

---

**Table S1.**
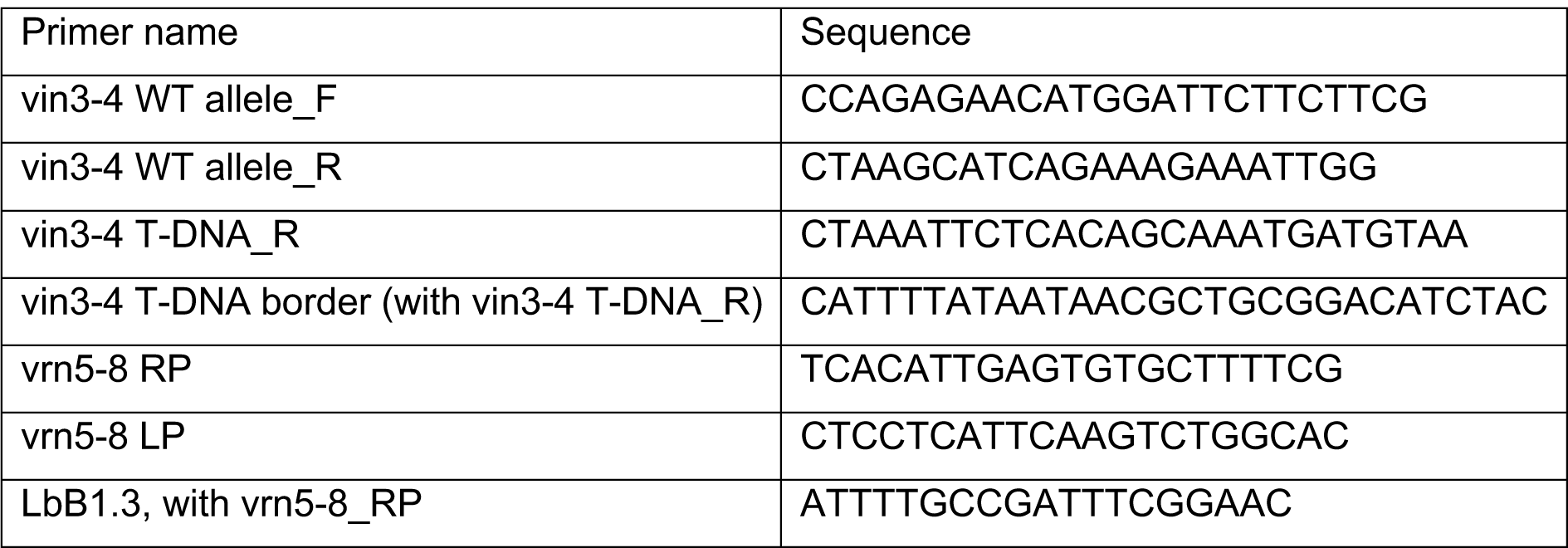
Primers used for generating the transgenic constructs

